# Hepatic *huntingtin* loss drives an acute phase response and liver injury in multiple mouse models

**DOI:** 10.64898/2026.07.28.741358

**Authors:** Colby L Samstag, Robert M Bragg, Kelsie Neumann, Ella Mathews, Kellie Lam, Christine C Wu, Deanna M Marchionini, Michael J MacCoss, Daniel Raftery, Jeffrey B Carroll

## Abstract

Multiple therapeutic strategies are being developed to slow Huntington’s disease (HD) progression through targeted reduction of huntingtin (HTT) protein or mRNA. Despite HTT’s discovery over 30 years ago, its cellular functions remain incompletely understood, and the long-term consequences of HTT-lowering therapies remain unclear. We previously demonstrated that hepatic HTT loss in mice disrupts hepatocyte zonation and metabolism. Here, we investigate the physiological consequences of hepatic *Htt* loss. Across multiple models of *Htt* loss—including ubiquitous and hepatocyte-specific genetic knockouts and a therapeutically relevant *Htt*-targeting siRNA—there was elevated expression of IL-6/STAT3-driven acute phase response genes. Single-nucleus RNA sequencing reveals a zonal pattern of hepatocyte stress, most highly upregulated in pericentral hepatocytes, and identifies a distinct pericentral cluster of stressed hepatocytes that was enriched ∼9.6-fold following *Htt* knockout. Histological examination reveals that *Htt* loss results in increased hepatic pathology, including hepatic intranuclear inclusions, apoptosis, and necrosis, as well as prevalence of granulomas. Transcriptomic analysis reveals significant upregulation of metallothionein genes following *Htt* loss, as confirmed by elevated plasma metallothionein-1 (MT1) levels in knockout mice. These findings underscore important safety considerations for HTT-lowering therapies and suggest candidate biomarkers for monitoring hepatic off-target effects in clinical trials.

## Introduction

Huntington’s disease (HD) is a devastating neurodegenerative disease caused by an expansion of a CAG triplet repeat in the huntingtin (*HTT*) gene (1). HD is an autosomal dominantly inherited disease, with a median age of onset at 40 years and is currently incurable. Despite the *HTT* locus first being mapped over three decades ago, the endogenous function of the protein remains imprecisely defined. HTT is widely expressed throughout the body with very little tissue-or cell-type specificity, and it has been implicated in diverse cellular processes including autophagy, microtubule transport, mitochondrial energy production, synaptic vesicle transport, and extracellular vesicle formation amongst others(2).

Accordingly, *HTT* is an essential gene: homozygous knockout of the mouse ortholog (*Htt*) is embryonic lethal by E8.5 (3–5), and complete loss of HTT is presumed incompatible with life in humans as well; no individuals with homozygous complete loss-of-function mutations have been identified(6). Rare living individuals have been identified carrying compound heterozygous hypomorphic mutations in *HTT*, a condition referred to as Lopes-Maciel-Rodan Syndrome (LOMARS) (6, 7). Affected individuals present with psychomotor developmental delay, hypokinetic movements, dystonia, and severe intellectual disability, although, unlike HD, these symptoms are neurodevelopmental and not progressive in nature. Notably, the parents of individuals with LOMARS are obligate carriers of partial loss of function alleles but have no apparent clinical presentation, suggesting that haploinsufficiency of *HTT* is well tolerated but that reduction below a threshold becomes pathogenic. The precise level of HTT that is required to sustain development remains unclear, as is the threshold required to maintain normal cellular function in adults, both of which have serious implications for HD therapeutics currently in development.

Due to its devastating nature, there is a tremendous push to develop therapeutics to modify the course of HD. New therapeutic targets have been discovered recently using genetic modifier and mechanistic studies which show great promise, yet the principal target for most HD drugs currently being developed is reduction of the HTT protein itself(8). Some of these therapeutics lower *HTT* expression specifically in the CNS; AMT-130 (UniQure) is delivered by neurosurgical infusion while tominersen (Ionis) is given by intrathecal injection, both of which spare peripheral organs but are invasive and do not lower *HTT* outside of the brain. Conversely, oral *HTT*-lowering medications— including votoplam (PTC518, PTC Therapeutics and Novartis AG) and SKY-0515 (Skyhawk Therapeutics)—offer a simpler route of administration and lower levels of *HTT* systemically. Notably, of the *HTT*-lowering programs currently in clinical-stage investigation, only three are predicted to be either allele-preferential (VO659 (Vico Therapeutics) and Takeda zinc finger repressors) or allele-selective (WVE-003, Wave Life Sciences) with all other programs lowering total HTT. Given the critical functions of the HTT protein, this strategy presents a challenge: identifying a safe therapeutic window, balancing the need to reduce levels of the mutant HTT protein sufficiently to spare the cell of its toxic effects while allowing enough HTT protein to fulfill its critical cellular functions. Indeed, inducible *Htt* knockout in adult mice causes a progressive and fatal neurodegenerative disorder with progressive sub-cortical calcification and a robust and rapid increase in plasma neurofilament light chain, indicating that HTT is required for neuronal health and its loss is neurotoxic (9, 10).

In addition to its well-established role in neuronal health, there is growing evidence that HTT is critical for healthy liver function. Transgenic mouse models of HD show altered gene expression of transcription factor master regulators of metabolic enzymes (11), while sheep models show significant alterations to fatty acid metabolism in the liver (12). In studies of people with HD (PwHD), several liver function tests are progressively abnormal and correlate with disease severity, demonstrating a subclinical but measurable decline in liver health as a feature of HD progression(13). Because the overwhelming majority of human HD work focuses on the brain, these phenotypes and their clinical significance remain largely unexplored. This is particularly relevant given that several orally administered *HTT*-lowering drugs are currently in development (e.g. Novartis [NCT07326709] and Skyhawk’s [NCT06873334]), with doses formulated to provide therapeutic levels of knockdown in the brain. Skyhawk’s candidate drug is reported to reduce *HTT* mRNA levels by 72% in the peripheral blood of healthy volunteers (8); given the liver’s central role in first-pass metabolism, hepatic HTT knockdown likely exceeds these levels. It is therefore imperative to study the endogenous function of HTT in the liver, as well as unintended consequences that may arise from HTT knockdown.

To better understand the impact of hepatic HTT lowering we previously characterized phenotypes in a hepatocyte-selective inducible *Htt* knockout mouse, termed Liver Knock Out (LKO)(14). We show that loss of *Htt* results in alterations to hepatocyte identity, changes to the zonation patterns of hepatocytes, altered expression of toxin catabolizing enzymes, and disruption of hepatic cell adhesion to Glisson’s capsule. Hepatocytes are typically arranged in a highly stereotyped fashion, with specialized metabolic functions matching their localization relative to the portal vein and the nutrients it provides(15). Bulk RNA sequencing revealed upregulation of genes associated with periportal cell identity and downregulation of genes associated with pericentral functions, liver identity genes, and hepatic transcription factors (14). Histologically, the highly stereotyped hepatic zonation patterns established and maintained by Wnt/β-catenin gradient were disrupted. We noted upregulation of immune related gene sets, though histological analysis revealed no appreciable differences in the frank immune cell infiltration between LKO and control animals, and only modest increases in the levels of circulating immune markers. Whether these immune signatures reflect a cell autonomous change consistent with transcriptional derepression, or whether they reflect activation of resident liver immune cells remains unresolved(14).

In this work we further explored the pathologic consequences of *Htt* reduction in the liver using complementary *in vivo* systems. Reanalysis of our existing RNAseq data using gene set enrichment analysis suggested that mice lacking hepatic *Htt* showed robust upregulation of inflammatory genes. To understand the physiological consequences of these gene expression changes, we utilized single-nucleus RNA sequencing to identify which cells underlie bulk gene expression changes and confirm the pathological consequences of these gene expression changes using additional histological profiling of liver tissue from *Htt* loss-of-function models. We also expanded on our findings by examining models of ubiquitous knockout, as well as a therapeutically relevant model of graded lowering using *Htt*-targeted siRNA. Our analyses show that *Htt*-lowering across all models leads to convergent transcriptional programs, including upregulation of metallothioneins, activation of acute phase and stress responses, and collagen remodeling. We conclude that both *Htt* knockout and robust *Htt* lowering are toxic to hepatocytes and causes immune cell activation within the liver.

## Results

### Liver-specific knockout of Htt in early development leads to inflammatory liver injury

To resolve the origin of immune gene upregulation in LKO livers, we first performed a careful reanalysis of our published transcriptomic data. We previously identified aberrant liver expression of genes normally enriched in other organ systems, including thymus and spleen(14). Reanalysis of these data revealed that these “tissue enriched” genes were predominantly immune-related genes, likely reflecting an aberrant liver immune response rather than a *bona fide* change to hepatic cellular identity (Supplemental Fig S1). To gain greater insight into the specific immune programs upregulated in LKO livers, we also performed Gene Set Enrichment analysis (GSEA) on this transcriptomic dataset. GSEA largely recapitulated our earlier EnrichR analyses, showing a profound reduction of oxidative phosphorylation and lipid metabolism genes (Fig 1A), but also revealed specific upregulation in genes from the interferon-gamma response, allograft rejection, TNFα signaling via NF-κB, and IL6/JAK/STAT3 signaling pathways (Fig 1A).

**Fig. 1:**
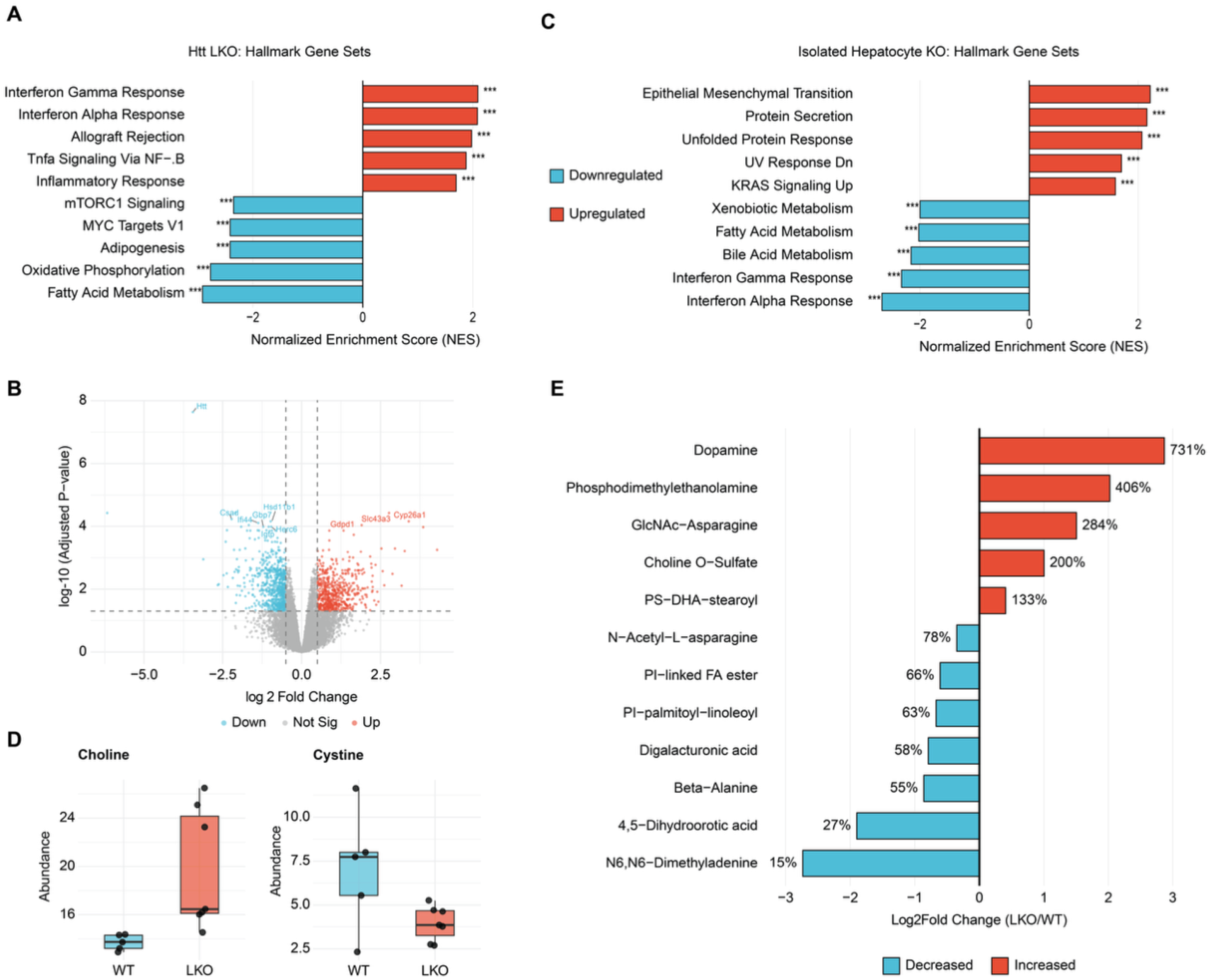
Developmental knockout of *Htt* in hepatocytes causes systemic metabolic disruption and immune gene upregulation. **(A)** Gene set enrichment analysis (GSEA) of differentially expressed genes in LKO versus control livers. Top 5 up-and downregulated pathways from the Hallmark gene set collection are depicted, as ranked by normalized enrichment score (NES). *padj < 0.05, ** padj < 0.01, *** padj < 0.001. Significance determined by fgsea’s multilevel Monte Carlo procedure with Benjamini-Hochberg adjustment **(B)** Volcano plot depicting differential gene expression in cultured hepatocytes isolated from LKO and control mice. Plot depicts log₂ fold change against - log₁₀(adjusted p-value), with genes meeting significance thresholds (|log₂FC| > 0.5, adjusted p < 0.05) colored if significantly upregulated (red) or downregulated (blue), or non-significant (gray). Dashed lines indicate fold-change and significance cutoffs. Of 15,726 genes analyzed, 685 were downregulated and 587 upregulated (|log₂ fold change| > 0.5, adjusted P < 0.05) **(C)** GSEA of cultured LKO versus control hepatocytes. Top 5 up-and downregulated pathways from the Hallmark gene set collection are depicted, as ranked by normalized enrichment score (NES). *padj < 0.05, ** padj < 0.01, *** padj < 0.001. Significance determined by fgsea’s multilevel Monte Carlo procedure with Benjamini-Hochberg adjustment **(D)** Plasma abundance of choline and cystine in 6-month-old LKO and Control mice (n=6/genotype) from targeted aqueous metabolite profiling. Significance determined by Student’s t-test. **(E)** Curated biological metabolites with significantly altered abundance in untargeted metabolomics of whole liver tissue from LKO versus control mice. Bars show log₂ fold change (LKO/Control). Significance determined by Welch’s t-test with Benjamini-Hochberg FDR correction. *padj < 0.05, **padj < 0.01, ***padj < 0.001

Our existing bulk liver RNAseq datasets cannot distinguish between immune cell infiltration and cell autonomous up-regulation of immune response genes in hepatocytes, so we isolated and cultured hepatocytes from LKO and control animals and performed RNA sequencing (Fig 1B). Gene Set Enrichment analysis on genes differentially expressed in LKO and control hepatocytes confirmed the dramatic reduction in metabolic programs, including fatty acid and bile acid metabolism, in LKO hepatocytes (Fig 1C). Isolated LKO hepatocytes displayed significant downregulation of immune genes (Fig 1C), indicating that *Htt* loss does not cell-autonomously de-repress immune gene expression in hepatocytes, and that the induction seen in vivo requires signals from the hepatic microenvironment that are absent in isolated culture.

To better understand the organismal consequences of the immune and metabolic changes, we next performed metabolomic profiling on *Htt* knockout animals and controls. We first performed targeted aqueous metabolomics on plasma from these animals to assess whether metabolic defects were detectable in circulation as a potential biomarker of hepatic *Htt* loss. This analysis revealed a significant increase in plasma choline (44% relative to control, nominal p = 0.025) and a decrease in cystine (44% reduction, nominal p = 0.045), but these did not survive corrections for multiple comparisons (Fig 1D). We also tested whether transcriptomic alterations were accompanied by changes in metabolites within LKO livers by performing untargeted global metabolomics on bulk liver tissue from LKO and control animals. This analysis revealed 12 significantly altered metabolites (FDR q < 0.05) (Fig 1E). LKO livers showed evidence of altered choline metabolism, with elevated levels of two choline-related metabolites, choline-O-sulfate (200% increase, q = 0.040) and phosphodimethylethanolamine (406% increase, q = 0.027), supporting the nominal increase in plasma choline. We also observed changes in metabolites associated with phospholipid and membrane composition, including decreases in two phosphatidylinositol species and an increase in phosphatidylserine-DHA-stearoyl (133%, q = 0.049). Taken together, these data indicate that hepatic *Htt* loss perturbs liver metabolism, particularly choline and phospholipid pathways, and that these changes are reflected in circulating metabolites.

### Multiple models of Huntingtin knockdown share a gene expression signature consistent with acute stress and upregulation of immune response

One of the principal motivations for our study is to identify potential off-target safety signals that might arise from HTT-lowering therapeutics. Our data suggest that hepatic loss of *Htt* was sufficient to lead to systemic metabolic and inflammatory changes, but it is impossible to differentiate between developmental defects versus the acute effects of *Htt* loss from the LKO model alone. In order to differentiate between these effects and to interrogate the relevance of physiological lowering of huntingtin in adult PwHD, we next performed RNAseq on livers from two previously published mouse models of inducible *Htt* reduction: 1) our “conditional knock out” (cKO) model which develops with normal *Htt* levels but undergoes ubiquitous excision of *Htt* after administration of tamoxifen at two months of age (10) and 2) mice treated with pan-*Htt* targeting siRNA (HTT10150)(16) conjugated to N-acetylgalactosamine (GalNAc), a ligand taken up specifically by hepatocytes via the asialoglycoprotein receptor(17). Livers were extracted from these mice and subjected to bulk RNA sequencing, in tandem with age-and sex-matched controls (see Materials and Methods section for additional details). We ran differential expression analysis for each model, comparing the transcriptome from cKO and siRNA-treated animals with their respective controls. (Supplemental Fig S2). As expected, both models showed significant reduction in hepatic *Htt* (cKO: 87% reduction, adjusted p = 6.8 × 10⁻⁴⁹; siRNA: 60% reduction, adjusted p = 1.7 × 10⁻² in bulk liver tissues). Because we were interested in identifying consistent pathways perturbed across models, we looked at concordant gene expression changes across all three models: LKO, cKO and siRNA-treated animals versus their respective controls (Fig 2A; a full list of treatments and animals used in this study can be found in Supp Table 1).

**Fig. 2.**
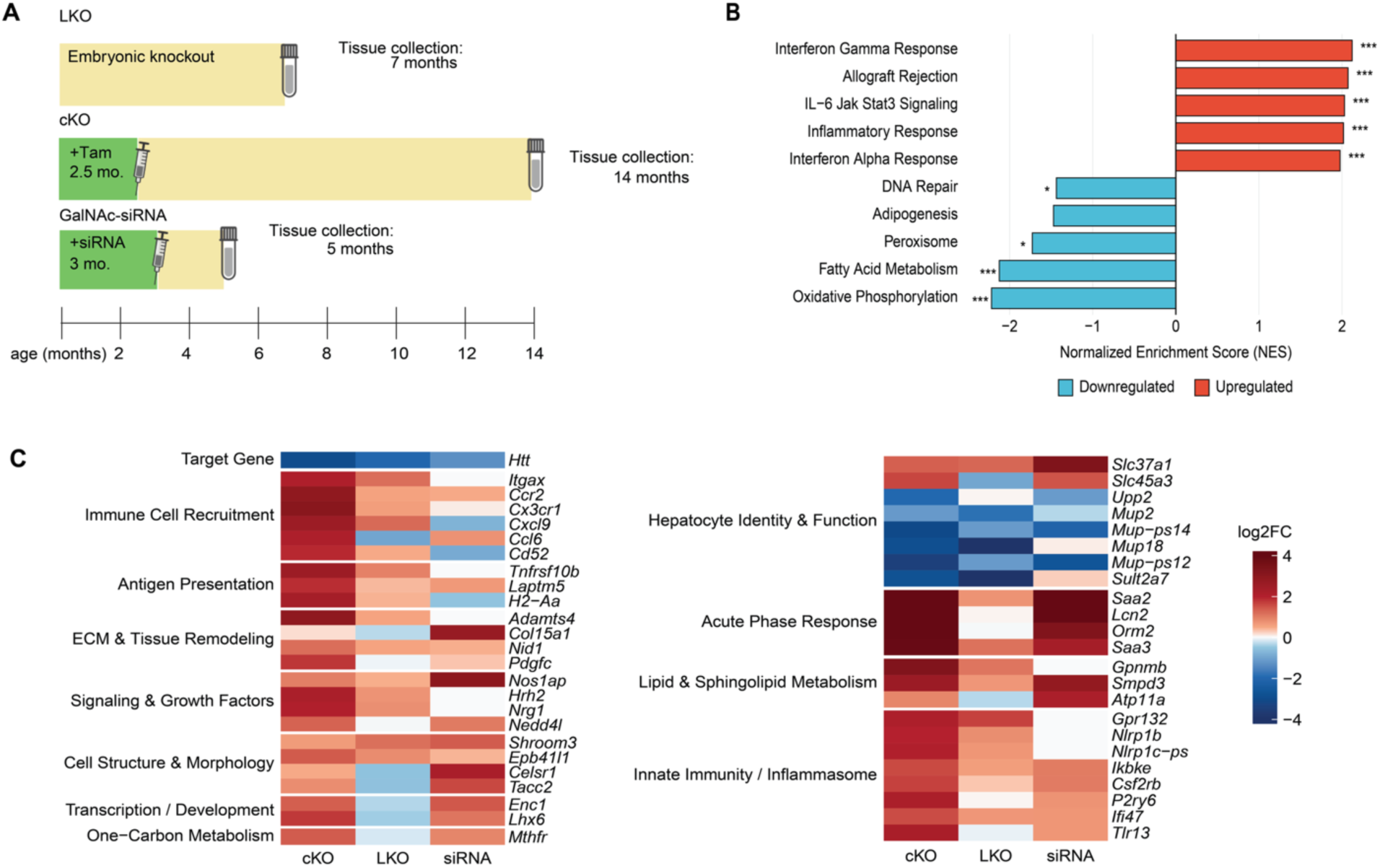
Multiple models of *Htt* lowering share convergent upregulation of inflammatory and immune genes in bulk liver tissues. **(A)** Schematic of timing of *Htt* loss and liver tissue collection of the mice used for comparative RNAseq analysis **(B)** Gene set enrichment analysis (GSEA) of combined differential gene expression across LKO, cKO, and siRNA models. Genes present in all three datasets with concordant direction of log₂ fold change were retained, and gene-level statistics were combined using Fisher’s combined method with GSEA performed on the resulting combined ranking. The top 5 up-and downregulated pathways against the MSigDB Hallmark gene set collection are depicted, as ranked by normalized enrichment score (NES). *padj < 0.05, ** padj < 0.01, *** padj < 0.001. Significance determined by fgsea’s multilevel Monte Carlo procedure with Benjamini-Hochberg adjustment. **(C)** Heatmap of genes with significant concordant gene expression changes following *Htt* knockdown, determined by Fisher’s combined p-value. Color scale represents log₂ fold-change relative to respective controls, with genes arranged by functional group

To determine whether shared cellular pathways were perturbed following *Htt* loss, we applied Fisher’s method to combine p-values across the full gene expression matrix, identifying genes and pathways with consistent directional signal. We ran GSEA on the combined differential gene expression and discovered pathway alterations that were largely concordant with our analysis of LKO alone, suggesting *Htt* loss results in similar downstream consequences across modalities. The pathways most highly upregulated in our combined dataset included the Interferon Gamma and Interferon Alpha pathways and the IL-6/JAK-STAT3 pathways (Fig 2B). Bile acid metabolism, amino acid metabolism, and electron transport chain genes were among the most consistently downregulated. Together, these suggest that the metabolic and inflammatory transcriptional changes in LKO animals were not the result of developmental liver injury but instead represented a common response to HTT lowering.

To better understand the cellular mechanisms underlying these pathway-level changes, we also identified which specific genes were most consistently differentially regulated across *Htt* loss models using Fisher’s combined statistic. Few genes were consistently downregulated across conditions, but those that were predominantly included several Major Urinary Proteins, a family of rodent-specific secreted lipocalins that regulate glucose and lipid metabolism (Fig 2C)(18, 19). Although MUPs lack a direct human homolog, their downregulation across *Htt* loss models point to conserved disruption of hepatic lipid and glucose metabolism(20). There was also consistent downregulation of *Sult2a7*, a member of the hepatic phase II detoxification sulfotransferase *Sult2a* family(21), as well as the pyrimidine salvage enzyme *Upp2*(22), both consistent with broad hepatic metabolic dysfunction. Conversely, upregulated genes include the sphingolipid metabolism gene *Smpd3* (23) and the one-carbon metabolism enzyme *Mthfr* (24), suggesting broad lipid remodeling following *Htt* loss.

Despite differences in mouse models and the timing and delivery of HTT lowering, our data revealed that inflammatory and immune markers were consistently upregulated across models (Fig 2C). Upregulated genes included pattern recognition receptors (*Nlrp1b*, *Tlr13*)(25), interferon-responsive genes (*Ifi47, Ikbke*) (26, 27), myeloid activation and cytokine signaling components (*Csf2rb, Gpr132, P2ry6*)(28–30), and the MHC II gene *H2-Aa*(31), consistent with activation of both innate and adaptive immunity responses. We also observed upregulation of extracellular matrix remodeling genes (*Adamts4, Col15a1, Nid1*) (Fig 2C)(32, 33) and the activated macrophage marker *Gpnmb,* consistent with stellate and Kupffer cell activation. Acute phase inflammation genes, typically expressed by hepatocytes in liver injury, were also upregulated across lowering models (*Saa2*, *Saa3*, *Orm2*, *Lcn2*)(34–36). We interrogated these datasets for evidence of shared upstream cellular mechanisms, including the unfolded protein response, hypoxia, oxidative stress, and macroautophagy, but found little consistent support for a single unifying mechanism across all three models (Supplemental Figure S3). Taken together, our RNAseq data suggest that loss of *Htt*, regardless of modality, leads to transcriptional changes that include disruption of metabolism and upregulation of acute phase inflammatory and fibrosis genes.

### Single-nucleus RNA sequencing shows robust hepatic upregulation of metallothioneins in LKO

Our data suggest that multiple modalities of *Htt* lowering trigger hepatic injury and an innate immune response, but bulk transcriptomics cannot resolve the nature of this injury, nor whether this toxicity affects all hepatocytes or is limited to a particular subset. Our previous data suggest hepatocytes shift towards a “periportal-like” metabolic reprogramming but does not differentiate whether this is due to a loss of hepatic programming, a gain of periportal-function, or loss of pericentral hepatocyte function. To resolve this, we performed single nucleus RNA sequencing on livers extracted from LKO animals and controls. Using the 10x Genomics single nucleus gel emulsion platform with 3’ capture, we recovered a total of 28,000 isolated nuclei across both conditions. After performing stringent quality control for mitochondrial and ribosomal RNA content, transcript diversity, and multiplet removal, we recovered a total of 24,471 high-quality nuclei (N=12,088 control, 12,383 LKO) with a median of 6,353 UMIs/nucleus. After performing standard snRNAseq analysis pipelines, our data recapitulated the major cell types of the liver, including hepatocytes, stellate cells, LSECs, Kupffer Cells, T/NK Cells, B Cells, and Erythroid cells (Fig 3A), as confirmed by mapping established marker gene expression (Supplemental Figure S4).

**Fig. 3.**
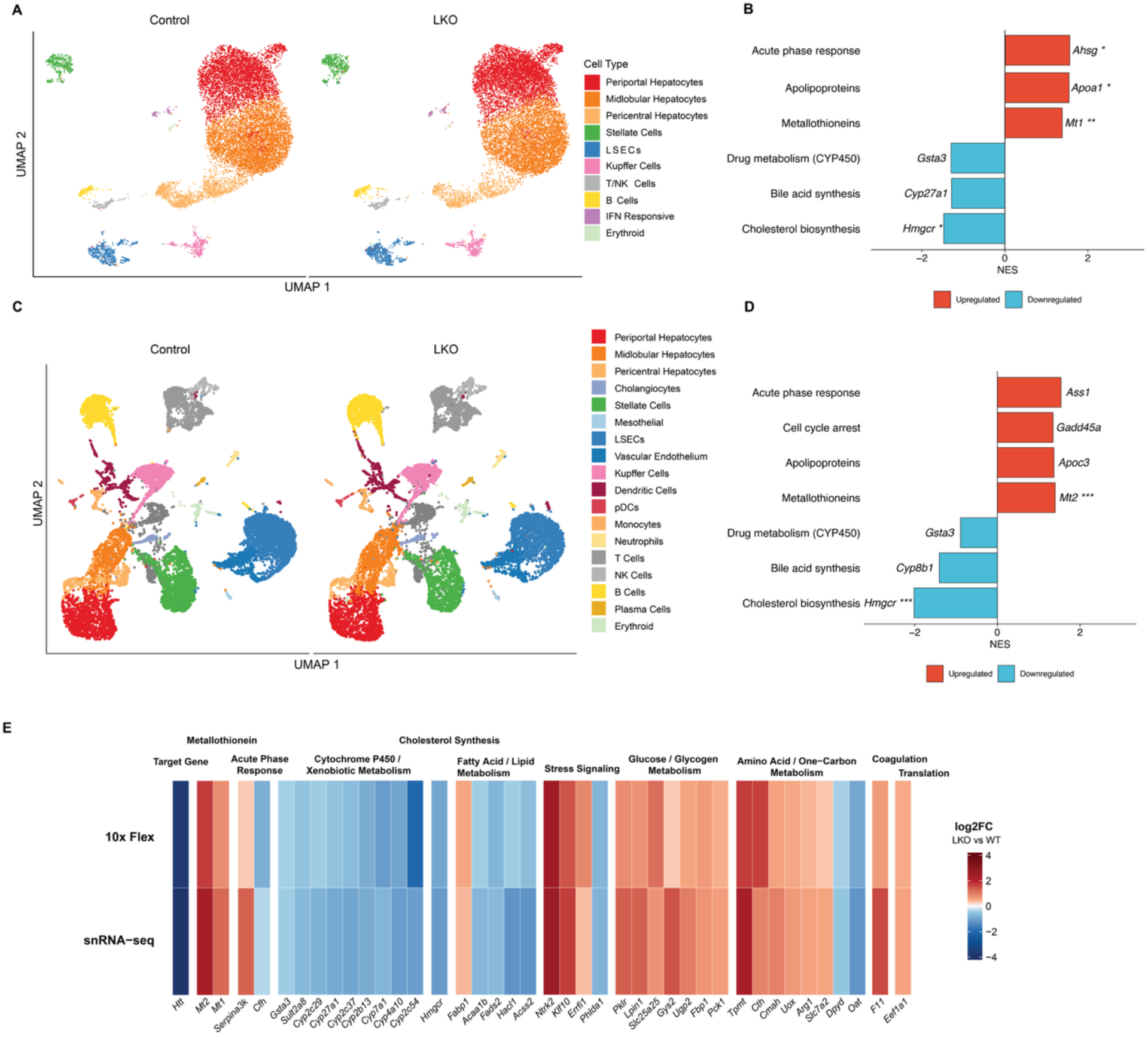
Single-nucleus and single-cell RNA sequencing of LKO livers reveals hepatic induction of acute phase and metallothionein genes. **(A)** UMAP visualization of 24,471 liver nuclei showing major liver cell types in control and LKO samples. **(B)** Differential gene expression in LKO versus control hepatocytes of pathways previously identified in bulk RNAseq. A full list of genes for these annotations can be found in Supplemental Table 2. Bars represent the normalized enrichment score (NES) for each pathway from fgsea, run on genes ranked by the Wilcoxon test statistic in LKO versus control hepatocytes. For each pathway, the gene with the largest absolute log2 fold change is labeled. **(C)** UMAP visualization of 26,678 liver cells identifying major liver cell types from scRNAseq of control and LKO samples. **(D)** Differential gene expression in LKO versus control hepatocytes of pathways previously identified in bulk RNAseq. Bars represent the normalized enrichment score (NES) for each pathway from fgsea, run on genes ranked by the Wilcoxon test statistic in LKO versus control hepatocytes. For each pathway, the gene with the largest absolute log2 fold change is labeled. (**E)** Heatmap of selected genes with concordant gene expression changes in LKO hepatocytes versus control in both snRNAseq and scRNAseq experiments, grouped by functional category. Color scale depicts log₂ fold change (LKO/Control) with values >=±4 clipped to preserve dynamic range. Genes depicted were among those reaching statistical significance (Benjamini-Hochberg adjusted *P* < 0.05) and |log₂ fold change| > 0.25 in both platforms in the same direction of change.

Mapping of hepatic zonal markers revealed a clear gradient from pericentral to periportal hepatocytes, confirming the utility of our sequencing results to capture hepatic diversity (Figure 3A). There was an apparent trend towards reduction in the relative proportion of pericentral hepatocytes and a corresponding increase in periportal hepatocytes in LKO animals compared to controls, but this could not be tested given the small sample size (Supplemental Figure S5). Our data also confirmed efficient hepatic *Htt* knockout: *Htt* was detected in 29.2% of control hepatocytes and only 0.6% of LKO hepatocytes. We next performed differential gene expression analysis, restricting our analysis to only hepatocyte clusters. We next tested whether pathways identified through our bulk RNAseq experiments reflected changes in hepatocytic expression by comparing average expression of these genes in LKO and control hepatocytes. This revealed coordinated downregulation of cholesterol biosynthesis (including *Hmgcr*, *Fdft1*, *Nsdhl*, *Lss*, and *Msmo1*), bile acid synthesis and transport (*Cyp7a1*, *Cyp27a1*, and *Abcb11*), and drug metabolism CYP450 genes (including *Cyp2b10* and *Cyp2a5*), alongside upregulation of acute phase response genes (*Serpina3* family, *Hp*, *Apcs*) and apolipoproteins (*Apoa1*, *Apoc1-3*, *Apoe*) in LKO hepatocytes, recapitulating the changes observed in bulk data (Fig 3B). We also noted that LKO hepatocytes had upregulation of pro-apoptotic *Bcl2l11* alongside reduced anti-apoptotic *Bcl2l1*, though the apoptosis gene set did not show coordinated enrichment. Notably, we discovered very significant upregulation of the metallothionein genes, which encode cytoplasmic cysteine-rich metal-binding proteins. Among the top differentially expressed genes, *Mt1* (avg log₂FC = 1.23, adjusted P < 2.2 x 10⁻³⁰⁸) was detected in 80.3% of LKO vs. 57.0% of control hepatocytes and *Mt2* (avg log₂FC = 2.33, adjusted P < 2.2 × 10⁻³^00^) was detected in 28.6% vs. 7.1% hepatocytes.

Capture of hepatocytes for snRNAseq is complicated by their fragility and by the abundance of cytoplasmic enzymes released during nuclei dissociation(37). To complement our snRNAseq experiment, we also performed single-cell RNAseq (scRNAseq) using the 10x Flex probe-based assay on formalin-fixed tissues on LKO and control livers. 26,678 cells were recovered following QC (N=13,975 control, 12,703 LKO). Hepatocytes represented a smaller fraction of the total pool of cells in scRNAseq compared to snRNAseq (∼21.5% of all cells N=5,736 hepatocytes), potentially reflecting greater capture of non-parenchymal and immune cells in the Flex assay, as well as the fact that multinucleate cells are overcounted in snRNAseq.

We investigated the same pathways as in snRNAseq and found largely concordant results, including upregulation of metallothioneins (*Mt1*, *Mt2*) and acute phase response genes (*Saa1*, *Saa2*, *Lcn2*), and downregulation of cholesterol biosynthesis and bile acid synthesis enzymes (*Cyp8b1*, *Cyp7a1*, *Cyp27a1*) (Fig 3D). Drug/xenobiotic metabolism genes showed mixed changes in the Flex dataset, with downregulation of *Cyp2e1* and *Cyp1a2* but upregulation of *Ugt2b37*. To identify hepatocyte expression changes robust to technical differences between these assays, we also analyzed the top 50 genes by absolute log2FC expression with concordant direction in snRNAseq and scRNAseq. This again revealed consistent downregulation of lipid and xenobiotic metabolic genes and upregulation of metallothioneins and stress response genes in LKO hepatocytes (Fig 3E). This analysis also revealed upregulation of glucose and one carbon/amino acid metabolic genes in LKO hepatocytes (Fig 3E). Taken together, these data support that chronic, hepatocyte-specific loss of *Htt* induces widespread metabolic changes and measurable hepatic stress.

### Single nucleus RNAseq following adult Htt loss reveals a subpopulation of hepatocytes with acute phase stress response induction

We next investigated whether inflammatory changes are induced in hepatocytes following acute loss of *Htt* in adult animals. To test this, we performed snRNAseq on a model of ubiquitous adult knockout, the cHtt-cQ20 model. The cHtt-cQ20 model was generated through insertion of loxP sites flanking exon 1 in the Q20 knock-in mouse model (Marchionini et al., in preparation). When crossed to UBC-Cre^ERT2^ mice, the resulting HttQ20^2lox/2lox^;UBC^cre/+^ progeny (termed *cQ20* hereafter) develop with normal *Htt* before ubiquitous excision following administration of tamoxifen, resulting in a 71% decrease in HTT protein in liver at 16 weeks of age compared to untreated animals.

snRNAseq of cQ20 liver samples yielded 43,174 high-quality nuclei passing QC (N=20,126 control, N=23,048 cQ20) comprising 13 cell types (Fig 4A). A high proportion of recovered nuclei were again derived from hepatocytes (∼79.1% of all nuclei), and mapping of canonical liver zonation markers revealed a clear gradient of periportal, midlobular, and pericentral hepatocytes (Supplemental Figure S4). We discovered a very modest shift toward increased periportal hepatocytes and decreased pericentral hepatocytes in cQ20 relative to control animals (Supplemental Figure S5). When analyzed for differential gene expression relative to control hepatocytes, cQ20 hepatocytes showed coordinated downregulation of cholesterol biosynthesis (*Nsdhl*, *Msmo1*, *Hmgcr*, *Idi1*, *Fdft1*), bile acid synthesis (*Cyp7b1*, *Cyp7a1*), and drug/xenobiotic metabolism genes (*Ugt2b1*, *Cyp1a2*, *Gstp1*) (Fig 4B). Like LKO hepatocytes, cQ20 hepatocytes showed upregulation of acute phase response genes (Fig 4C), but cQ20 hepatocytes additionally showed strong upregulation of cell cycle arrest genes (*Cdkn1a*, *Btg2*, *Gadd45a/g*, *Trp53inp1*) and the pro-apoptotic *Bcl2l11*. There was also much greater induction of hepatic metallothioneins in cQ20 than in LKO: *Mt1* was increased by 6.80 log₂FC (expressed in 86.4% of hepatocytes in cQ20 vs. 5.2% in controls) and *Mt2* by 7.21 log₂FC (71.8% of cQ20 hepatocytes expressing vs. 1.7% in controls).

**Fig 4.**
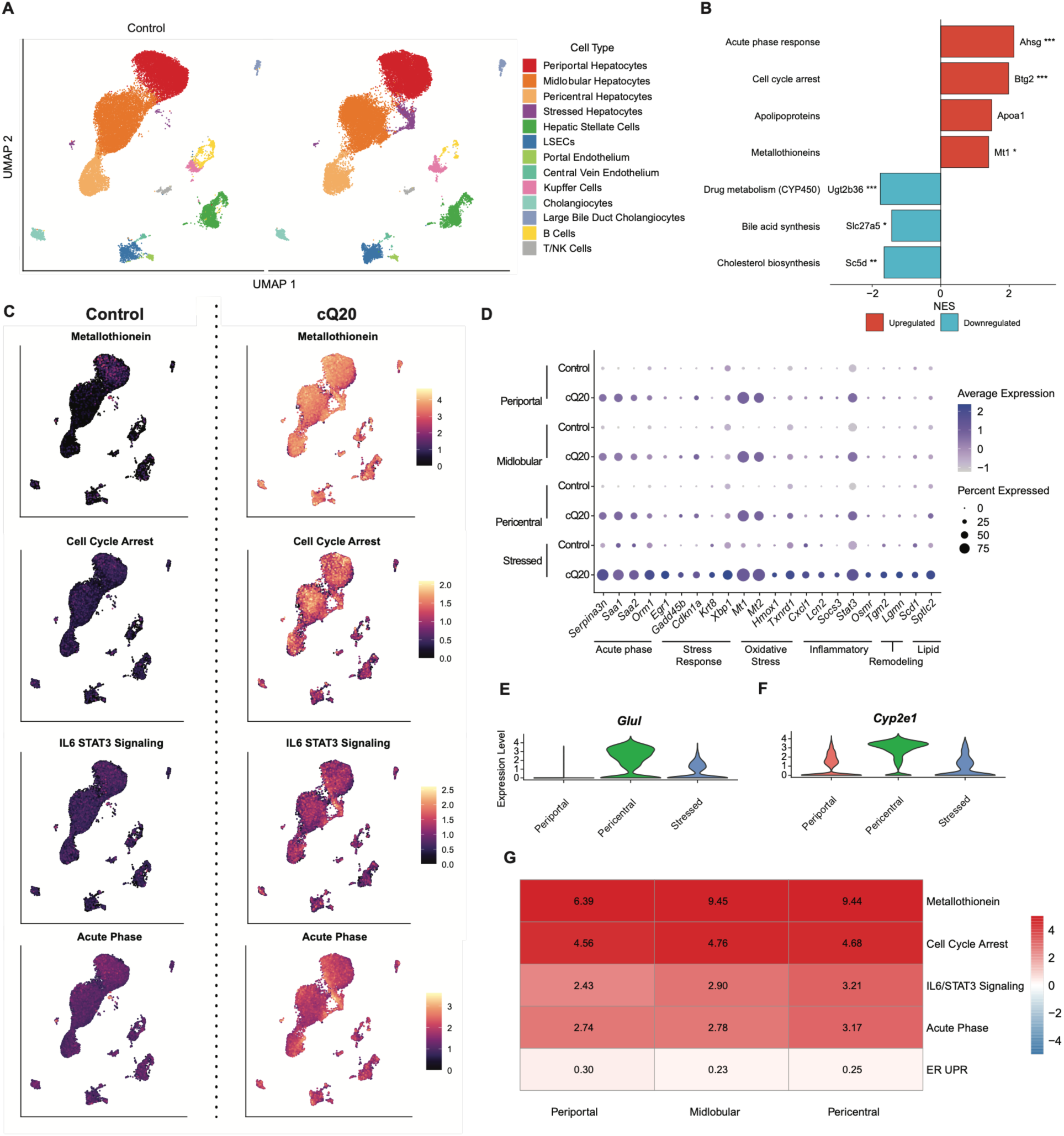
Single-nucleus RNA sequencing of cQ20 and control livers reveals a novel cluster of stressed hepatocytes and pericentral enrichment of acute stress response genes following adult *Htt* loss. **(A)** UMAP visualization of 43,174 hepatic nuclei showing major liver cell types in control and cQ20 samples. **(B)** Differential gene expression in cQ20 versus control hepatocytes of pathways previously identified in bulk RNA-seq. A full list of genes for these annotations can be found in Supplemental Table 2. Bars represent the normalized enrichment score (NES) for each pathway from fgsea, run on genes ranked by the Wilcoxon test statistic in cQ20 versus control hepatocytes for each pathway, the gene with the largest absolute log2 fold change is labeled. **(C)** UMAP projections colored by the mean log-normalized expression of genes from the indicated pathways, including metallothioneins (*Mt1* and *Mt2*; top), cell cycle arrest genes (second), IL6/STAT3 family members (third), and acute phase response genes (bottom) in Control and cQ20 livers. Color scale indicates mean log-normalized expression per nucleus. **(D)** Dotplot depicting expression of marker genes with significantly higher expression in the stressed hepatocyte cluster. Expression is stratified by cluster and genotype, with the color representing the Z-scaled expression of each gene and the size of the dot representing the proportion of cells in each cluster expressing that gene (non-zero counts). **(E, F)** Expression (in counts) of (E) *Glul* and (F) *Cyp2e1* from nuclei derived from pericentral, periportal, and stressed hepatocyte snRNAseq clusters. **(G)** Heatmap of stress-response signature induction across the portal–central hepatocyte axis following adult *Htt* loss. For each hepatic zone (periportal, midlobular, pericentral), nuclei were aggregated into pseudobulk profiles by animal, and differential expression (cQ20 vs. control) was computed per zone using DESeq2. Each cell shows the mean per-gene log₂ fold-change (maximum-likelihood estimate) across the detected genes in a signature. Genes expressed in fewer than 5% of hepatocytes were excluded. Color represents log₂ fold-change (cQ20/control), with the values printed in each cell. Signatures shown are metallothioneins, acute phase response, IL-6/STAT3 signaling, cell cycle arrest, and the ER/unfolded protein response, with gene lists provided in Supplemental Table 3. The stressed hepatocyte subcluster was excluded due to insufficient representation in control samples.

Unexpectedly, we also discovered a unique subset of hepatocytes abundant in cQ20 animals but almost entirely absent in control animals (see circled cluster in Fig 4A). We performed stringent quality control checks and normalization algorithms to confirm that this cluster did not exist solely as the result of technical artifacts, and we also confirmed that it did not differ significantly in QC metrics when compared to other clusters. This unique cluster persisted through all iterations of QC and clustering parameters, verifying that it is derived from a true biological cell population rather than a technical artifact. Both genotypes have nuclei that map to this cluster, but they were markedly enriched in cQ20 livers: this cluster comprises 0.51% of hepatic nuclei in control animals, but 4.9% of hepatic nuclei in cQ20, a 9.6-fold increase. This cluster was defined by high expression of many of the same genes that were upregulated in our bulk RNAseq datasets following *Htt* loss, including acute phase response genes (*Saa1*, *Saa2*, *Serpina3n*, *Fgb*, *Fga*, *Fgg*, *Orm1*, *Hp*, *Hpx*, *Lbp*), stress-responsive transcription factors (*Junb*, *Jun*, *Egr1*, *Xbp1*, *Atf4*, *Atf6*, *Hif1a*), metallothioneins (*Mt1*, *Mt2*) and redox/antioxidant regeneration genes (*Hmox1*, *Txnrd1*, *Glrx*, *Gsr*). The cluster also showed induction of NF-κB pathway members (*Nfkb1*, *Nfkbib*, *Rela*, *Ripk1*, *Tifa*, *Tnip2, Bcl3*), structural cytokeratins associated with hepatocyte injury (*Krt8*, *Krt18*), and markers of cell cycle arrest and apoptosis (*Cdkn1a*, *Bcl2l11*, *Mcl1*, *Trp53inp1*). This cluster also showed upregulation of IL-6/JAK-STAT signaling components (*Stat3*, *Socs3*, *Socs2*, *Jak1*, *Jak2*, *Il6st*, *Osmr*) (38). Collectively, this transcriptional profile is consistent with a hepatocyte state undergoing an IL-6/STAT3-driven acute phase response coupled with acute stress, recapitulating features from our bulk RNA sequencing of *Htt* lowering. We term this cluster “stressed hepatocytes.” Immune and stress response genes broadly upregulated in cQ20 hepatocytes reach their highest expression in the stressed hepatocyte cluster (Fig 4C). When stratified by cell type and genotype, these genes show the greatest increases in both per-cell expression levels and the proportion of expressing cells in the cQ20 stressed hepatocyte cluster (Fig 4D).

From this finding, we hypothesized that pericentral hepatocyte dysfunction or death could explain the shift towards a more periportal-like state that we previously observed in LKO animals. To test this hypothesis, we first looked at markers of pericentral cell identity between clusters. *Glul* (glutamine synthetase) is one of the most exquisitely sensitive markers of pericentral hepatocytes. An enzyme that detoxifies ammonia into glutamine, *Glul* expression is restricted to the 2-3 layers of hepatocytes directly surrounding the central vein(15). In our clustering, there was high expression of *Glul* in the pericentral hepatocyte cluster but virtually no expression in the periportal cluster, suggesting clear separation of nuclei with minimal contamination (Fig 4E). There was also high expression of *Glul* in stressed hepatocytes, consistent with their being pericentral in origin (Fig 4E). We also looked at levels of *Cyp2e1*, which marks pericentral hepatocytes and contributes to alcohol and xenobiotic detoxification. *Cyp2e1* was highly expressed in pericentral hepatocytes, low in periportal hepatocytes, and intermediate in stressed hepatocytes (Fig 4F). The retention of *Glul* and reduction of *Cyp2e1* in stressed hepatocytes is consistent with their downregulation of metabolic function, possibly in response to acute stress.

Finally, we compared expression of stress response pathway genes in the portal-central hepatocyte axis, comparing cQ20 expression to control in each hepatocyte compartment. The very low number of stressed hepatocytes in control animals precludes the comparison of stressed hepatocyte clusters directly. However, in the remaining hepatocyte clusters there was a gradient of upregulation of stress response pathways, with pericentral hepatocytes showing the greatest upregulation and periportal the least, particularly in acute phase response genes (Fig 4G). Together, these analyses reveal a pericentral hepatocyte population vulnerable to *Htt* loss, with a stress characterized by acute phase induction, IL-6/STAT3 signaling, cell cycle arrest, and upregulation of metallothioneins.

### Histological analysis reveals hepatotoxicity and immune upregulation following Htt loss

Our gene expression and metabolomic data indicate shifts in the metabolic capacity of hepatocytes in tandem with the upregulation of inflammatory genes. To determine whether these molecular changes manifested as gross pathological changes in the liver, we evaluated livers of LKO and cKO mice for several markers of gross pathology. Firstly, we revisited whether there was an increase in frank immune cell infiltration following *Htt* reduction. In our previous work, we did not see an apparent increase in the number of immune cells in LKO mice (14). Given the upregulation of immune markers across RNAseq datasets, we revisited this in LKO and extended this analysis to cKO samples. Assessment of portal immune cell infiltration showed no significant differences in LKO or cKO from their respective controls, though several knockout animals were noted to have mild infiltration (Supp Fig S6). We next evaluated samples for the presence of granulomas, organized collections of immune cells that form in response to chronic inflammation or foreign material and arise from coordinated activation of resident macrophages and lymphocytes (39). Pathological assessment revealed no difference in the presence of granulomas between LKO animals and controls (Fig 5B), but a significant increase in granuloma formation in cKO animals (Fig 5C). Consistent with the upregulation of both adaptive and innate immune pathways seen in RNAseq, the increased prevalence of granulomas indicates that *Htt* loss leads to local inflammatory lesions but absence of widespread immune cell infiltration.

**Fig. 5.**
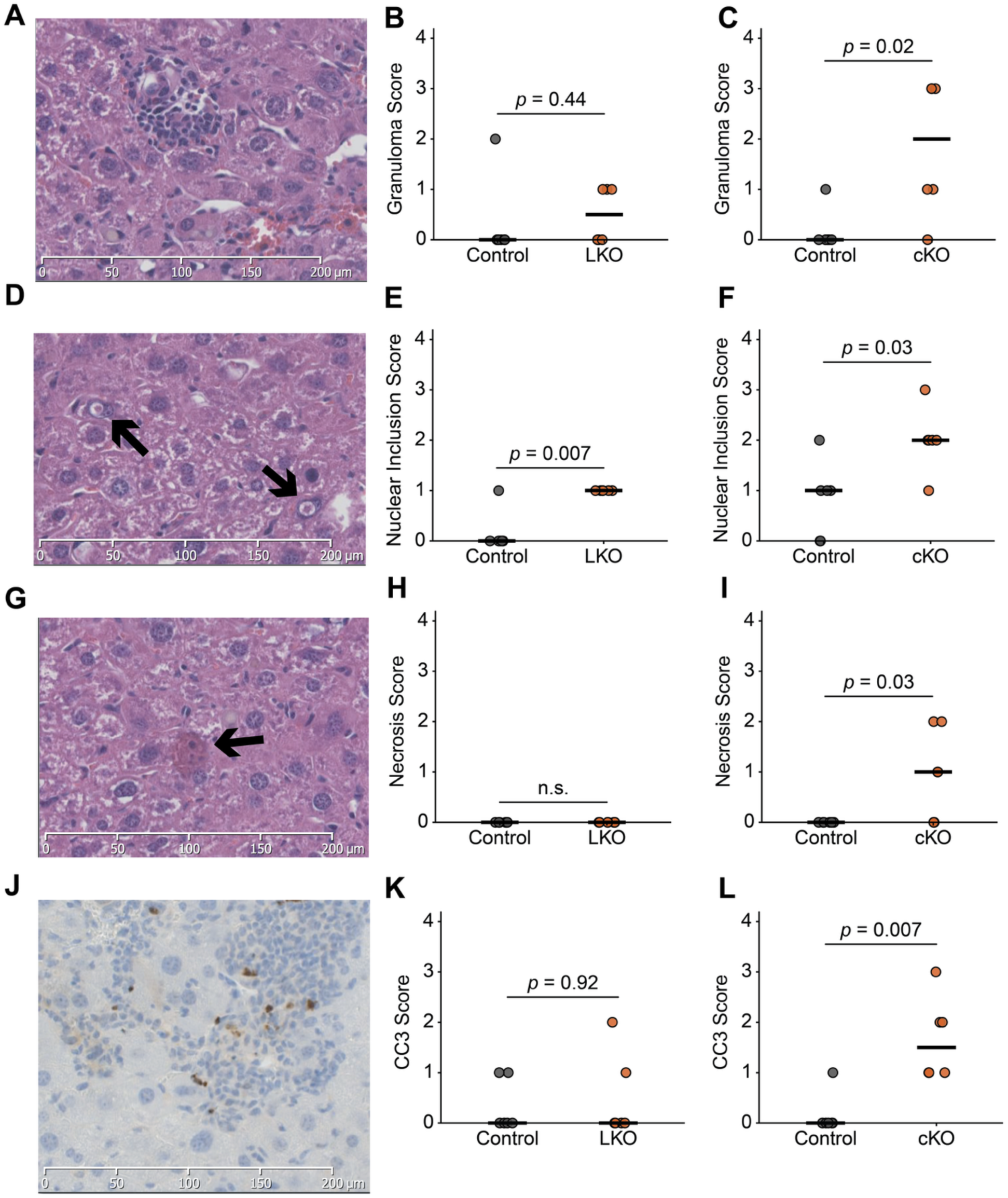
Histology reveals gross hepatic pathologic changes after *Htt* knockout. **(A)** Representative H&E-stained slide depicting granuloma in cKO liver section. Pathological assessment of granuloma score for **(B)** LKO and **(C)** cKO**. (D)** Representative H&E-stained slide depicting intranuclear inclusion in cKO liver section. Pathological assessment of intranuclear inclusion score for **(E)** LKO and **(F)** cKO. **(G)** Representative H&E-stained slide depicting hepatic necrosis in cKO liver section. Pathological assessment of necrosis score for **(H)** LKO and **(I)** cKO. **(J)** Representative Cleaved caspase-3 immunostained slide counterstained with hematoxylin depicting CC3+ cells and nuclei. Pathological assessment of CC3 score for **(K)** LKO and **(L)** cKO. For each semi-quantitative histological score animals were blindly scored on a 0-4 scale, and groups were compared using the Wilcoxon rank-sum test. Each point represents one animal. P-values are shown for comparison of knockout versus respective control.

We next evaluated whether the changes in hepatocyte gene expression and metabolism led to detectable hepatic pathology. We evaluated samples for the presence of intranuclear inclusions, aggregates of ubiquitinated and misfolded proteins that accumulate under conditions of proteotoxic, metabolic, and oxidative stress, and increase with liver injury and aging (40–42). Both LKO and cKO animals displayed a significant increase in intranuclear inclusions relative to their controls (Fig 5D-F). We also evaluated the extent of hepatic necrosis and apoptosis, using H&E staining and cleaved-caspase 3 immunostaining, respectively. There was no significant difference in either apoptosis or necrosis between LKO and controls (Figs 5H, 5K), but there was a significant increase in both pathologies in cKO mice (Fig 5I, 5L). While the pathological features of *Htt* loss were significantly increased in cKO knockout animals when compared to their controls, they remained sparse within the tissue despite the high efficiency of *Htt* knockout. In tandem with the findings from snRNAseq, these findings suggest that most hepatocytes tolerate Huntingtin loss while a subset of predominantly pericentral hepatocytes are vulnerable to hepatotoxicity.

### Circulating markers of hepatic injury following Htt loss

Our transcriptomic, metabolomic, and histologic analyses together show that *Htt* loss leads to a hepatic acute stress response, localized immune activation, and metabolic rewiring. To identify potential risks for clinical reduction of *HTT*, we also searched for circulating biomarkers that predict hepatotoxicity in the plasma of mouse models following *Htt* lowering. For this, we leveraged two complementary approaches: an unbiased proteomics approach and candidate gene approach informed by our transcriptomic analyses.

To determine whether proteomics could detect novel secreted proteins following *Htt* loss, we collected plasma from 7mo cKO mice and controls, enriched for extracellular vesicles, and analyzed this fraction by mass spectrometry (Fig 6A). We previously demonstrated that this EV fractionation technique efficiently reduces bias from the most highly expressed plasma proteins and more accurately reflects the cargo from cells of origin, allowing for more sensitive detection of biomarkers(43). Gene set enrichment analysis of differentially abundant proteins revealed significant upregulation of extracellular matrix organization and glycoprotein biosynthesis pathways, and significant downregulation of proteins associated with mRNA processing and mitochondrial translation (Fig 6B). Notably, the proteins most enriched in cKO mice included cathepsin G, proteinase 3, and neutrophil elastase, all serine proteases released upon neutrophil degranulation and none of which were detected from our bulk liver RNA sequencing datasets. These data demonstrate that *Htt* loss leads to detectable inflammatory and immune signatures in the plasma but could reflect broad tissue injury and neutrophil activation rather than a specific signature of hepatocyte stress. To search for candidate biomarkers specific to hepatocyte stress, we returned to our transcriptomic datasets.

**Fig. 6.**
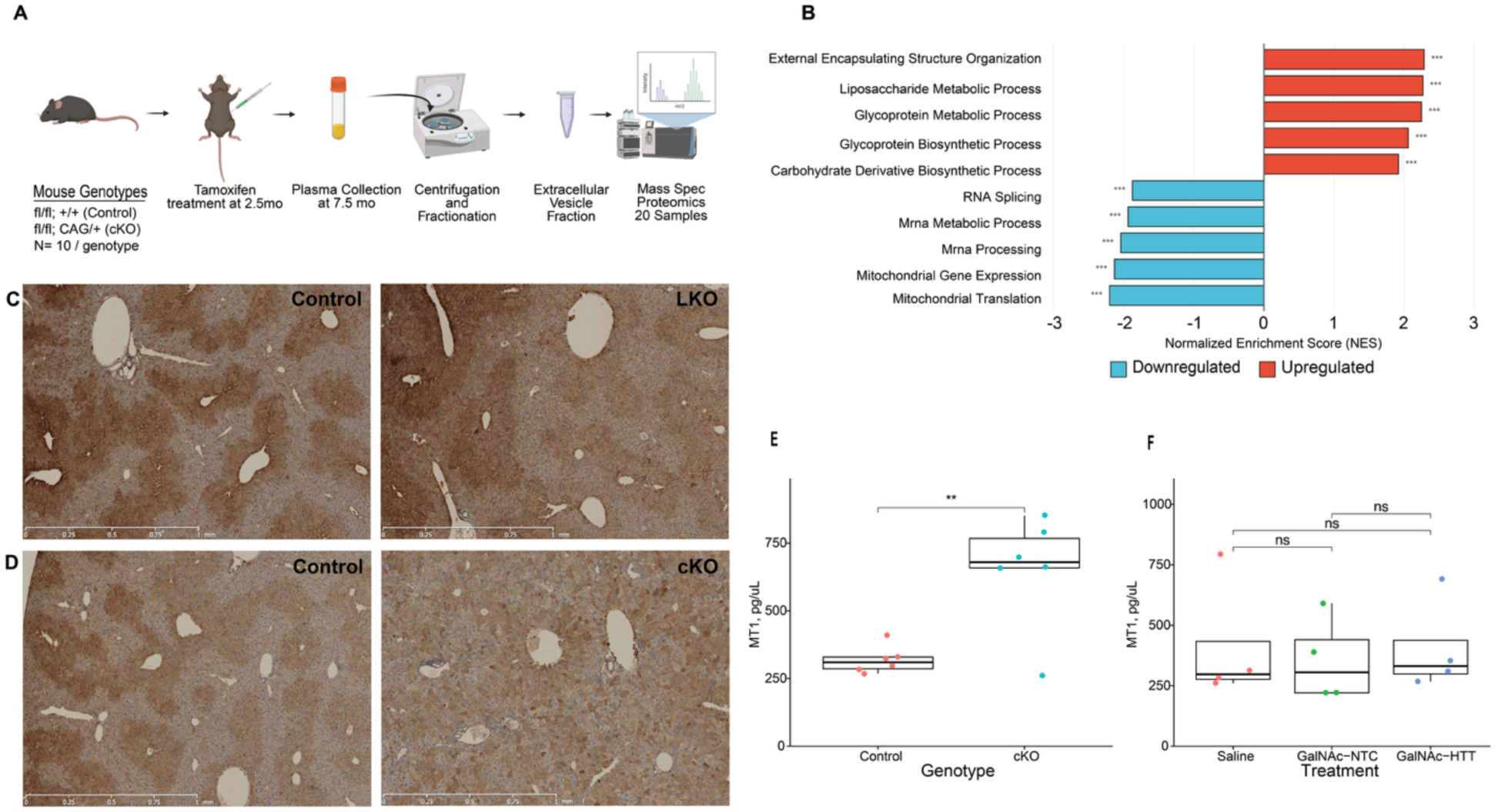
Plasma biomarkers reveal hepatic stress following *Htt* loss. **(A)** A schematic of our plasma EV proteomics workflow, including mouse genotypes, treatment, and mass spec library prep. **(B)** GSEA (GO: Biological Process) of differentially abundant proteins in plasma EVs between cKO and controls. Pathways ranked by NES, top five pathways up-and down-regulated shown. *padj < 0.05, ** padj < 0.01, *** padj < 0.001. Representative MT2 immunohistochemistry in liver sections from **(C)** LKO and **(D)** cKO animals and their respective controls. Plasma concentrations of metallothionein-1 from **(E)** cKO and **(F)** HTT siRNA-treated animals as determined by ELISA. Significance for panel B determined by fgsea’s multilevel Monte Carlo procedure with Benjamini-Hochberg adjustment. Significance for panels E and F determined by two-tailed Welch’s t-test.

In searching for candidate biomarkers, we prioritized three criteria: consistent upregulation across multiple models of *Htt* loss, enrichment in the stressed hepatocyte cluster identified in cQ20, and supporting evidence of being detectable in plasma. The most promising candidate genes that emerged from this analysis were the metallothioneins, *Mt1* and *Mt2*, which are best characterized for their role in maintaining cellular metal ion homeostasis and scavenging reactive oxygen species during acute oxidative stress and heavy metal exposure. Metallothioneins emerged repeatedly across our snRNAseq datasets—*Mt1* was increased 2.35-fold and *Mt2* was increased 5.03-fold in LKO hepatocytes compared to control hepatocytes in snRNAseq. Metallothionein upregulation was far more dramatic in cQ20 animals—112-fold and 148-fold increase in hepatocytes for *Mt1* and *Mt2* respectively. Furthermore, the subpopulation of stressed hepatocytes showed even greater enrichment (182-fold increase in *Mt1* expression and 240-fold increase in *Mt2* expression over control hepatocytes), consistent with its induction in the cell-autonomous response to *Htt* loss. Finally, multiple studies document alterations to plasma metallothionein levels in wide-ranging liver pathologies (44–47), supporting their potential as circulating biomarkers of hepatocyte stress.

To confirm whether transcriptomic alterations are reflected in protein-level changes to metallothioneins, we performed immunohistochemistry on liver sections from cKO and LKO animals and their respective controls, probing for metallothionein 2 (Fig. 6C, D). Staining confirmed that hepatocytes are the major cellular source of metallothionein 2 in the liver. In control animals, expression followed a classic zonal pattern, with high pericentral staining and markedly reduced signal in periportal and midlobular regions. This pattern was disrupted in knockout animals, with more pronounced disruptions in cKO livers, consistent with their more severe pathological changes in histology (Fig. 6C, D). While our IHC was not suited to quantify changes in the total abundance of metallothionein 2, these data nonetheless confirm the pericentral hepatic expression of metallothionein and demonstrate the disruption of this expression pattern following *Htt* knockout.

Finally, we tested whether the dysregulation of metallothioneins observed in the liver was reflected in circulation. We compared plasma MT1 levels from cKO and *Htt* siRNA-treated animals along with their respective controls using ELISA. Plasma MT1 levels in cKO animals were significantly higher than in controls, demonstrating that liver injury in these animals is sufficient to change circulating levels of MT1 (Fig. 6E). However, MT1 levels were not significantly different between siRNA-treated animals and controls (Fig. 6F), consistent with its milder transcriptional dysregulation. Together, these data identify metallothionein as a putative noninvasive plasma biomarker that tracks with the severity of liver injury following *Htt* reduction.

## Discussion

*HTT* lowering remains the predominant therapeutic strategy for HD, yet most programs in development reduce expression of both the wild-type and expanded alleles systemically. For many of these drugs, hepatocytes will likely experience some of the highest levels of *HTT* reduction. Here, we observe that robust *Htt* lowering is associated with clear hepatotoxic sequelae. Our findings demonstrate that even partial loss of HTT function induces acute hepatic stress, underscoring the need to carefully define the therapeutic window for systemic *HTT*-lowering strategies.

Our previous work demonstrated transcriptional upregulation of the liver’s immune response following complete *Htt* loss but left several questions unanswered: whether adult loss or reduction of *Htt* recapitulated these gene expression changes, and where these changes originate given the apparent lack of circulating immune cells in plasma or in infiltrating immune cells in the liver (14). Our current work addresses these gaps. Across multiple knockout and acute *Htt*-lowering models, we find convergent immune upregulation including hepatic IL-6/JAK-STAT3 signaling, collagen remodeling, and altered metabolic programming. The absence of overt hepatic immune infiltration suggests a focal paracrine response, consistent with Kupffer cell-derived IL-6 activating a cytoprotective STAT3 program in hepatocytes at the site of injury(48). All our models of *Htt* lowering showed upregulation of STAT3 target genes (*Saa1, Saa2, Saa3, Socs3*), with substantially greater induction in models of adult-onset *Htt* loss than in developmental loss, despite comparable reductions in the HTT protein. Additionally, snRNAseq of cQ20 livers reveals that *Htt* knockout induces a subpopulation of stressed hepatocytes of pericentral origin, enriched for transcription of acute phase genes, consistent with previous studies showing that stressed hepatocytes can also be a source of paracrine IL-6 signaling(48, 49). Together, these data suggest that adult *Htt* knockout is less well tolerated than embryonic knockout, perhaps due to a greater capacity for compensatory mechanisms available during development. Given the liver pathological findings after adult *Htt* lowering, our data have direct implications for therapeutics in adult PwHD.

Our data also showed that *Htt* loss is associated with down-regulated pathways critical for normal hepatic function including cholesterol biosynthesis, bile acid synthesis, and xenobiotic metabolism. Bulk RNA seq on liver tissues from multiple models of *Htt* lowering show clear and consistent upregulation of acute inflammatory response pathways, IL-6/JAK-stat signaling, and induction of cell cycle arrest and apoptotic genes. snRNAseq analysis reinforced these results: both developmental and adult knockout of *Htt* resulted in a broad hepatic stress response consistent with bulk RNAseq, while adult knockdown leads to the development of a novel cluster of acutely stressed pericentral hepatocytes. Histology confirms the presence of increased hepatic apoptosis and necrosis, as well as granuloma formation, supporting the theory that *Htt* loss leads to focal hepatotoxicity, particularly in pericentral regions. Taken together, our work suggests that the metabolic changes that we had previously observed in the liver following *Htt* loss occurs concomitantly with an acute inflammatory response, though the causality of this relationship remains unclear.

The dramatic upregulation of metallothioneins in response to *Htt* loss was unexpected but is also consistent with an acute inflammatory response.

Metallothioneins are classically regarded as metal-responsive genes whose expression is tightly regulated by cellular levels of zinc and copper. However, the promoters of metallothioneins also have response elements for multiple STAT proteins (including STAT3), glucocorticoids, and specificity protein 1 (Sp1), all of which can independently induce metallothionein expression(50). Further, IL-6 administration alone is sufficient to rapidly and robustly induce metallothioneins(51). Metallothionein expression in *Htt* knockdown models closely mirrors the expression of acute phase proteins. In bulk RNAseq, *Mt1* and *Mt2* levels remained essentially unchanged in LKO animals but were highly upregulated in acute knockdown (2.3-and 1.9-fold increase in cKO, 5.6-and 13-fold increase in siRNA-treated animals). We see no evidence of induction of other metal stress response pathways, so their upregulation is best explained as a component of the acute phase response rather than a primary response to metal dyshomeostasis. Notably, metallothioneins are thought to play a protective role in HD. Early work showed that overexpression of metallothioneins including MT3 protects against polyQ aggregation and toxicity in yeast and mammalian HD cell models, though the mechanism was unclear(52). Recently, single-nucleus sequencing from postmortem PwHD samples identified a neuroprotective subpopulation of metallothionein-expressing reactive astrocytes(53). Follow-up studies found that a SNP in the metallothionein gene cluster is a genetic modifier in human HD age of onset and demonstrated *in vitro* that MT3-expressing astrocytes have increased glutamate buffering(54). Our findings suggest that MT1/2 may serve a protective role in hepatocytes after *Htt* loss analogous to the putative neuroprotective role of MT3 after expression of mutant HTT. If so, they could serve as a powerful biomarker for identifying hepatic stress resulting from HTT lowering in the liver.

The role of HTT in maintaining liver function in humans is only beginning to be understood, but mounting clinical evidence suggests progressive hepatic function decline in PwHD. For example, the ^13^C-methionine (^13^CO₂) breath test, a noninvasive measure of hepatic mitochondrial oxidative capacity, reveals subclinical impaired mitochondrial function even in people with premanifest HD (55) and tracks longitudinally with disease progression(55, 56). Although the molecular mechanisms linking HTT to hepatic function have primarily been studied in the mutant HTT context, recent work suggests substantial overlap of gene networks perturbed in both HD and HTT loss-of-function contexts (57). At the cellular level, hepatocytes share key steps of the pathogenic cascade that have come to define HD development in the central nervous system. As in the brain, the livers of HD mouse models show transcriptional dysregulation (58), mitochondrial dysregulation (59), defects in autophagy (60, 61), and somatic instability of *HTT’s* CAG tract(62, 63). Knock-in modified mouse models including R6/2(11) and HdhQ150(64) also show hepatic accumulation of mutant HTT-containing intranuclear inclusions, though to our knowledge these pathogenic features have not been demonstrated in liver samples from PwHD.

Although we found no consistent transcriptional signature of autophagy dysregulation across models (Supplemental Figure S3), *Htt* knockout models exhibit histologically confirmed increased intranuclear inclusions, a feature linked to autophagy dysfunction across a range of liver pathologies (65, 66). Recent work has demonstrated that hepatic *Htt* knockout impairs the autophagic flux of ApoE (60), which may explain the transcriptional disruption to cholesterol metabolism genes seen in *Htt* knockout mice. Both liver and plasma metabolomics also show that *Htt* loss resulted in disruptions to one-carbon and phospholipid metabolism, pathways whose disruption are well established drivers of hepatic steatosis and nonalcoholic fatty liver disease (67, 68). Taken together, our data suggest that *Htt* loss-of-function affects autophagy alongside cholesterol and phospholipid metabolism, with direct implications for HTT-lowering therapeutics.

One limitation of our study is our reliance on mouse models, as the mouse immune system and liver anatomy diverge considerably from those of the human. Further, because LKO and cQ20 snRNAseq datasets differ in animal age, nuclei isolation chemistry, and sequencing depth, comparisons of effect magnitude between these datasets are qualitative. Finally, limited tissue availability prevented us from testing every assay across all genotypes, ages, and sexes. A systematic analysis of the time course of pathology onset following HTT lowering was beyond the scope of this study but warrants future investigation.

In this work, we have shown convergent inflammatory phenotypes across multiple models of *Htt* loss, including a therapeutically relevant model of *Htt* lowering. Particularly for drugs that are expected to lower HTT ubiquitously, our data suggest that monitoring for signs of hepatic inflammation should be incorporated into the safety assessment of HTT-lowering clinical trials.

## Materials and Methods

### Sex as a Biological Variable

Our study examined male and female animals, and similar findings are reported for both sexes.

### Mouse models

Multiple strategies were used to generate HTT knock-out mouse lines and tissues, with all lines on a consistent B6 background and a full matrix of the animals, ages, and experiments is provided in Table S1. A hepatocyte-specific HTT KO was generated by crossing Albumin-cre mice (RRID: IMSR_JAX:003574) with HTT-flox mice to generate *Htt*^LKO/LKO^, as described previously(14, 69, 70). A whole-body HTT KO was generated by crossing a tamoxifen inducible UBC-cre (RRID:IMSR_JAX:007001) line with HTT-flox as described previously(10). A separate inducible whole-body HTT KO was generated by utilizing a CAG-cre in the same manner as the UBC-cre line, as previously described(10). HTT-KO was induced for these two lines by treating mice with tamoxifen (75 mg/kg) for 5 consecutive days at 2-mo of age. A third inducible whole-body HTT KO was generated by crossing the inducible UBC-cre line (RRID:IMSR_JAX:007001) with a distinct HTT-flox (Htt-Q20^2lox^; Marchionini et al, in preparation). Briefly, this flox line contained a humanized HTT exon 1 knock-in with a CAG-tract length of 20, a non-pathogenic length in humans, and exon 1 was flanked by LoxP sites. HTT-KO of this line was induced at 12 weeks of age by treating with tamoxifen (100 mg/kg) for 5 days at Jackson Laboratories, a fully AAALAC-accredited animal facility.

### GalNAc-siRNA-HTT lowering

Liver-specific HTT lowering was induced by treating C57Bl/6 (RRID:IMSR_JAX:000664) mice with GalNAc-conjugated siRNA as previously described (Trivalent GalNAc conjugated to HTT10150) (16, 71). Mice were IP injected with 10 mg/kg of siRNA at 3-months-old, again at 4-months, and were sacrificed at 5-months old.

All mice were fed ad libitum and kept on a standard 12/12 hour light/dark cycle. All experiments utilizing mice were approved by the appropriate IACUC committees. HttLKO and HTTcKO mice were part of UW animal protocol 4387-03. Portions of this work were conducted under contract at The Jackson Laboratory, an AAALAC-accredited facility in good standing. All animal-related methods and procedures were performed in accordance with The Jackson Laboratory’s IACUC-approved animal use summaries, established standard operating procedures, and the institution’s OLAW assurance.

### Tissue collection

Livers of LKO, cKO, and siRNA-treated mice were collected after overdose from IP injection of pentobarbital (Euthasol), flash frozen, and stored at -80°C until use. cQ20 mice were euthanized via cervical dislocation following the AVMA Guidelines for the Euthanasia of Animals (2020 edition), and livers processed similarly. For histology, mice were transcardially perfused with PBS followed by 10% neutral buffered formalin and livers were additionally fixed for 72hrs before being transferred to PBS. Livers were dehydrated, embedded in paraffin, and mounted on slides in 5 um sections. For experiments using purified hepatocytes, a 2-step collagenase perfusion was used to dissociate and collect frozen pellets or cultured for 24 hr on collagen-coated plates as described(72).

### Metabolomics

Aqueous metabolites were extracted from plasma (n=12) by methanol-based protein precipitation and from liver tissue (∼10 mg) by Dounce homogenization in aqueous methanol, as previously described(73). Stable isotope-labeled internal standards (SILISs) were added during extraction and reconstitution to monitor sample preparation and enable absolute quantification of 30 metabolites. Plasma was analyzed by targeted LC-MS in MRM mode on an AB Sciex 6500+ triple quadrupole coupled to Shimadzu UPLC, with HILIC separation on a Waters XBridge BEH Amide column and dual-polarity ESI (216–220 metabolites detected; median QC CV 5.5%). Liver extracts were analyzed by untargeted HPLC-ESI-MS on Agilent 6520 and 6545 Q-TOF instruments with HILIC separation on a Waters XBridge BEH Amide column and an ammonium acetate/acetic acid gradient. Data were processed in Progenesis QI (Waters) with putative identifications against HMDB (10 ppm tolerance). Liver data were normalized to BCA protein content. Additional methods can be found in Supplemental Note 1.

### Bulk RNAseq

RNAseq from bulk tissues and purified hepatocytes was completed as previously described at Azenta(14). Briefly, RNA was extracted using RNeasy Lipid Tissue Mini Kit (Qiagen). RNA with RIN >7 was carried forward to library prep, using the Illumina TruSeq kit according to the manufacturer’s instructions. cDNA libraries were sequenced using Illumina sequencers (2×150bp). Poor quality reads were removed and adaptor sequences trimmed from the resulting BAM files using Trimmomatic v.0.36. STAR aligner v.2.5.2b was used to align reads to the mouse reference genome (mm10), and count files were generated using Subread package v.1.5.2. Differential gene expression was compared between knockout animals and their respective controls using limma and edgeR.

Differential gene expression datasets from cKO, LKO, and siRNA-treated mice and their respective controls was used as input for gene set enrichment analysis (GSEA). For GSEA performed on individual datasets (LKO liver and isolated LKO hepatocytes), genes were ranked for differential expression according to sign(log₂FC) × −log₁₀(adjusted p-value). For consensus analysis, LKO, cKO, and siRNA treated liver datasets were combined using Fisher’s combined probability method, whereby combined p-value was calculated as χ² = −2 Σ ln(*pᵢ*), evaluated against a chi-squared distribution with 2*k* degrees of freedom, with *k* representing the number of datasets in which the transcript was observed. Genes were then ranked by sign(mean log₂FC) × −log₁₀(combined p-value). For both individual RNAseq datasets and the Fisher’s combined metric, GSEA was performed using the R package fgsea, using Hallmark, GO Biological Process, and Reactome gene set collections from MSigDB. For other graphical representations, the R packages ggplot2, ComplexHeatmap, pheatmap, and EnhancedVolcano were used. Claude (Sonnet and Opus 4 models, Anthropic, claude.ai) was used to assist with iterating computational code for data analysis and identifying candidate genes of interest. All AI-assisted outputs were reviewed, validated, and verified by the authors.

### snRNAseq libraries

Single-nucleus suspensions were generated from liver tissues stored at -80°C for LKO (14-month-old, n=2/genotype) and cQ20 (4-month-old, n=2/genotype). Suspensions were made for LKO using the Chromium Nuclei Isolation Kit (10x Genomics) according to the manufacturer’s protocol, and nuclei were counted on a hemocytometer following trypan blue exclusion staining. Tissues from cQ20 animals were shipped to Azenta (South Plainfield, NJ) on dry ice, and nuclei were isolated using the Nuclei Extraction Buffer (Miltenyi Biotec) with gentle MACS dissociation in C tubes according to the manufacturer’s protocol. Nuclei were counted using trypan blue exclusion on a Countess III automated cell counter (Thermo Fisher).

Single-nucleus RNAseq libraries were prepared using the Chromium Single Cell 3’ kit (10x Genomics) with a target capture of 8,000 gel bead-in-emulsions (GEMs) per sample for LKO experiments and 15,000 GEMs per sample for cQ20 experiments. Libraries were assessed for quality on an Agilent Bioanalyzer 2100 (LKO) or Agilent TapeStation (cQ20), quantified by Qubit 2.0 fluorometry (Invitrogen), and pooled libraries were quantified by qPCR (Applied Biosystems) prior to sequencing on an Illumina NovaSeq (LKO) or HiSeq 4000 (cQ20) to generate 2×150bp reads. Raw bcl files were demultiplexed and converted to FASTQ, sequence reads were aligned to the mouse reference genome (GRCm38) and quantified count matrices were generated using Cell Ranger (10x Genomics). To reduce the influence of ambient RNA contamination in LKO samples, FASTQ files were processed using Cell Ranger with the --force-cells=7000 parameter to retain only the top 7,000 nuclei per sample in downstream analyses.

### snRNAseq analysis

snRNAseq datasets were processed using a standard R pipeline. Briefly, filtered feature-barcode matrices for each sample were loaded into Seurat v5, excluding genes detected in fewer than 3 nuclei and nuclei with fewer than 200 detected features. Quality control metrics including the number of detected genes, total UMI counts, mitochondrial transcript percentage, and transcriptomic complexity (log10 genes per UMI) were computed for each nucleus. Nuclei were filtered to retain those with 200-6,000 (LKO) or 250-6,000 (cQ20) genes, 500-40,000 UMIs, mitochondrial percentage <5%, and complexity >0.8. Doublets were identified and removed using scDblFinder. Ambient RNA contamination was evaluated using SoupX, but estimated contamination was below the 2% threshold for all samples so ambient RNA correction was not applied. Per sample normalization was performed using SCTransform (v2). Principal component analysis was performed on the SCT assay, and batch effects corrected using Harmony. Nearest-neighbor graphs were constructed from 30 Harmony-corrected dimensions, clusters were identified using the Louvain algorithm at a resolution of 0.3, and UMAP dimensionality reduction was computed from the Harmony embedding. After initial clustering, clusters that were predominantly low-quality nuclei were identified by high expression of mitochondrial transcripts and ambient RNA markers (e.g., *Apoe, Apoa2, Hp*) and excluded from analysis, with re-processing using the same normalization, integration, and clustering parameters. From the revised clustering, cell types were annotated based on canonical marker gene expression, including zonally defined hepatocyte populations (*Glul, Cyp2e1, Sds, Hal*), Kupffer cells (*Clec4f*), liver sinusoidal endothelial cells (*Stab2, Ptprb*), stellate cells, cholangiocytes (*Pkhd1*), and immune cells (*Ptprc*). In the cQ20 dataset, a stressed hepatocyte population was identified based on high expression of acute phase response genes (*Serpina3n, Fgg, Fgb*). Differential gene expression was performed between knockouts and controls for each cell type using the Wilcoxon rank-sum test (min.pct = 0.1, logfc.threshold = 0.25). Additional plotting and visualization were done using ggplot2, EnhancedVolcano and patchwork. For the zonal gradient analysis, nuclei from periportal, midlobular, and pericentral hepatocytes were aggregated into pseudobulk profiles by animal and zone. Genes expressed in <5% of hepatocytes were excluded, and differential expression (cQ20 vs. control, n=2 animals/genotype) was computed per zone with DESeq2. Heatmap depicts the mean per-gene log₂ fold-change across each signature.

### Single-cell RNA sequencing (Flex)

Tissue stored at −80 °C was processed as follows. Briefly, for each sample 25 mg of liver tissues from LKO and control animals (n=2/genotype) were finely minced on dry ice and incubated in fixation buffer containing formalin (10x Genomics) at 4 °C for 20h. Samples were then processed according to the Chromium Fixed RNA Profiling (Flex) assay using Liberase TL tissue digestion. Suspensions were filtered through a 30 µm filter (Miltenyi Pre-Separation Filter) and counted by trypan blue exclusion. Transcripts were captured with the mouse Whole Transcriptome probe set (10x Genomics) according to the manufacturer’s protocol. Libraries were sequenced at Azenta on an Illumina NovaSeq X to generate 2 x 150bp reads, and probe-barcode count matrices were generated using Cell Ranger.

### Histological analysis

Immunohistochemistry on livers from 6-mo LKO animals and controls (N=6/genotype) and 14 mo cKO and their controls (N=6/genotype) was performed on formalin fixed paraffin-embedded sections according to protocols developed at the UW Histology and Imaging Core (RRID:SCR_028435). Antigen retrieval was completed using pH 6.0 Citrate buffer for 10 minutes (Metallothionein) and pH 9.0 EDTA buffer for 20 minutes (Cleaved caspase-3) in a decloaking chamber (Biocare, Pacheco, CA; catalog number DC2012) at 110°C. Following antigen retrieval, slides were blocked for 20 minutes with 5% Normal Goat Serum in PBS and incubated with primary antibodies antibodies for Cleaved caspase-3 (1:250 dilution; Cell Signaling; catalog number 9579S) or Metallothionein (1:500 dilution; Abcam; catalog number ab192385) for 30 minutes at room temperature. The primary antibodies were detected using the Bond automated stainer and Bond Refine Polymer Detection Kit (Leica Biosystems, DS9800). H&E-stained sections were scored for immune cell infiltration, intranuclear inclusions, granuloma prevalence, and hepatic necrosis by a veterinary pathologist blinded to sample group assignment. For CC3 staining, at least 5 fields of view at 20x magnification for each animal were scored by a trained laboratory technician blinded to sample group assignment. Each metric was evaluated on a predefined 0-4 semi-quantitative scale, with samples showing no pathology assigned a score of 0 and the maximum pathology a score of 4. The Wilcoxon-rank sum test was used to compare groups for each pathology measure.

### Plasma extracellular vesicle proteomics

Blood samples were collected from 7-mo cKO mice (N=10 /genotype) via cardiac puncture or submandibular vein puncture, balanced across genotypes, into K2EDTA tubes. Approximately 700 uL of cardiac blood was collected from each animal, centrifuged at 2,500g for 10min. The plasma fraction was snap frozen using dry ice and stored at -80°C. Extracellular vesicles were enriched from 25μL plasma using MagReSyn strong anion exchange (SAX) beads (ReSyn Biosciences) on a ThermoFisher KingFisher Apex as previously described (43), with a modification that trypsin digestion was performed in 50 mM Tris, pH 8.5. In parallel, 1 μL equivalent unfractionated plasma was digested using Protein Aggregation Capture using the same reduction, alkylation, and digestion conditions. For both preparations, 800 ng enolase standard was added as an internal quality control(74), and digestion was performed with Pierce porcine trypsin (ThermoFisher Scientific) at a ratio of 20:1 protein to trypsin for 1h at 47°C. Digests were quenched with 0.5% trifluoroacetic acid, spiked with Pierce Retention Time Calibrant (PRTC) peptide cocktail (ThermoFisher Scientific) to a final concentration of 50 fmol/μL, and stored at −80°C.

### Liquid chromatography–mass spectrometry

Peptide samples were analyzed by data-independent acquisition (DIA) on a Thermo Scientific Vanquish Neo UHPLC coupled to an Orbitrap Astral mass spectrometer with a heated CorSolutions source maintained at 40°C. Peptides were loaded on a Thermo Scientific PepMap Neo Trap Cartridge and separated on a Bruker PepSep C18 column (8 cm × 150 μm, 1.7 μm) attached to a 5 cm × 20 μm ID Sharp Singularity tapered emitter (Fossil Ion Tech) using a 21 min gradient from 4.5–38% acetonitrile in 0.1% formic acid at 1.0 μL/min. MS1 scans were acquired in the Orbitrap at 240,000 resolving power (m/z 200) over a scan range of m/z 400–900, followed by 167 DIA MS/MS scans on the Astral detector using 3 m/z non-staggered precursor isolation windows with window placement optimization with normalized collision energy of 27 assuming charge state 2 used for all MS/MS scans.

### ELISA

Plasma was collected from 7-month-old cKO (N=6 animals/genotype) via submandibular vein and 5-month-old siRNA treated (N=4 animals/genotype) via cardiac puncture and stored as above. Plasma metallothionein levels were measured using an ELISA kit for mouse MT1 (Novus Biologicals, cat. # NBP3-06945). Briefly, plasma was diluted 1:200 and incubated in a microplate coated with MT1 capture antibody according to the manufacturer’s instructions. All samples were run in duplicate and absorbance measured using a VICTOR Nivo (Revvity) plate reader.

## Supporting information

Supplemental Summary Data

## Data availability

All code used in our analyses is available via github repositories https://github.com/csamstag/liverKnockout (snRNAseq) and https://github.com/uw-maccosslab/collab-uw-carroll (proteomics). All transcriptomic datasets are available in GEO accession GSE329259. All proteomics data is available on Panorama.

## Acknowledgements

We thank Velvet Smith (University of Washington), Bereket Gidi (University of Washington), and Youjing Zheng (University of Washington) for help with histology analysis and figure preparation. We thank Jessica Snyder for her assistance with pathology scoring and histology experimental design, and Rotonya Carr (University of Washington) for consultations on liver anatomy and pathology. We thank all members of the Carroll Lab and Marie Y Davis for critical evaluation of this work.

## Author Contributions

CLS: conceptualization, software, data curation, formal analysis, supervision, investigation, visualization, project administration, and writing—original draft, review, and editing

RMB: writing—conceptualization, data curation, formal analysis, investigation, review and editing

KN: Investigation

EM: Investigation

KL: Investigation

CCW: Investigation

DM: conceptualization, resources, writing—review and editing

DR: Investigation, writing—review and editing

JBC: conceptualization, resources, formal analysis, supervision, funding acquisition, project administration, and writing—original draft, review, and editing

